# Transcription Start Site Heterogeneity Confounds the Landscape and Functional Interpretation of uORFs

**DOI:** 10.1101/2024.04.25.591216

**Authors:** Hsin-Yen Larry Wu, Qiaoyun Ai, Polly Yingshan Hsu

## Abstract

Upstream open reading frames (uORFs) are predicted in over half of the genes in many eukaryotes and often reduce translation and mRNA stability. This raises a paradox: why do uORFs remain so prevalent in eukaryotic genomes despite their repressive effects on gene expression? Using *Arabidopsis* as a model, we show that transcription start site (TSS) selection controls the inclusion and repressive impact of uORFs within the steady-state RNA pool. We find that TSS selection influences uORF inclusion in >90% of uORF-containing genes and selectively excludes uORFs from many transcripts. Although bioinformatic analyses predict uORFs in 55% of *Arabidopsis* genes, fewer than 10% of expressed transcripts contain them. This TSS filtering is nonrandom: genes whose uORFs are excluded are enriched for housekeeping functions, whereas genes retaining uORFs are associated with regulatory roles. We show that failing to account for TSS heterogeneity can mask length-dependent uORF repression and lead to misinterpretation of uORF activities. Importantly, uORFs mostly excluded from native mRNAs can still repress translation when artificially placed upstream of a reporter, demonstrating that apparent uORF repressiveness may not reflect their activity in native transcripts. Our findings establish TSS selection as a major determinant of uORF-mediated translational control. Because TSS heterogeneity is widespread among plant and animal genes, we warn that ignoring TSS heterogeneity can confound analyses of regulatory elements in mRNA 5′ leaders. We therefore recommend accounting for TSS usage when studying uORFs and other mRNA regulatory elements; otherwise, conclusions may reflect regulatory potential rather than regulatory activity in native transcripts.

**SIGNIFICANCE STATEMENT:** Upstream open reading frames (uORFs) are cis-regulatory elements in the 5′ leaders of eukaryotic mRNAs that suppress the translation of downstream main coding sequences. uORFs are predicted in more than half of genes in many eukaryotes. Their prevalence is paradoxical given their strong repressive potential. Analyzing transcription start site (TSS) usage in Arabidopsis, we show that uORFs are often excluded from major transcript isoforms because transcription initiates downstream of the element. However, conserved uORFs with established regulatory roles are preferentially retained. Thus, the scope of post-transcriptional regulation by uORFs is strongly shaped by transcriptional control. More broadly, ignoring TSS heterogeneity can lead to misinterpretation of the prevalence and regulatory impact of other mRNA features annotated in 5′ leaders.

## INTRODUCTION

Upstream Open Reading Frames (uORFs) are small ORFs located in the 5′ leaders or 5′ untranslated regions (5′ UTRs) of eukaryotic mRNAs. Genome-wide studies have shown that uORFs are widespread in multicellular eukaryotes, with bioinformatic analyses identifying uORFs in 64%, 60%, 60%, 55%, and 55% of genes in humans, mice, zebrafish, flies, and *Arabidopsis*, respectively (1). uORF translation can repress downstream main ORF (mORF, annotated ORF) translation and promote mRNA degradation, estimated to reduce protein levels by 30 to 80% (2–6). Given that uORFs exert strong repressive effects, it is puzzling that potential uORFs are widespread across diverse species. This "uORF paradox" raises the question of why cells would produce so many uORF-containing mRNAs despite their reduced translational output and the associated energetic cost.

An evolutionary framework proposes that uORFs follow two contrasting trajectories: beneficial uORFs are retained, whereas harmful uORFs are eliminated through purifying selection (7). The best-known examples of preserved uORFs are those encoding evolutionarily conserved peptides (CPuORFs) (8–15). In plants, approximately 120 CPuORFs have been identified and shown to sense metabolites, regulate abiotic and biotic stress responses, and control various aspects of plant physiology and development (16–23). Other uORFs that are extremely short or not conserved at the peptide-sequence level may still show conservation of uORF presence across species (24, 25). Conversely, several naturally occurring alleles that disrupt or weaken uORFs are associated with favorable agronomic traits. For example, *Arabidopsis* accessions lacking the *PHO1* uORF and a uORF SNP in soybean *GmPHF1* are linked to improved phosphate acquisition, variation in the rice *OsABA2* uORF influences preharvest sprouting, and population-scale uORF start-site variants in maize are associated with altered protein abundance and whole-plant phenotypes (26–29). Beyond plants, a similar balance between beneficial and detrimental uORFs is also evident in humans. For example, the uORFs of stress-responsive transcription factor *ATF4* enable stress-induced translational activation (30), whereas loss of a repressive uORF in the thrombopoietin gene causes hereditary thrombocythemia (31), and creation of a novel uORF in the tumor suppressor gene *CDKN2A* predisposes to melanoma (32). Together, these examples illustrate that uORFs can be either beneficial or detrimental and their inclusion or exclusion may be favored accordingly.

The inclusion of uORFs within mRNAs could potentially be regulated at the transcription level through changes in transcription start sites (TSSs), or at the post-transcriptional level through alternative splicing. Yet, the prevalence of either type of regulation on uORFs is largely unknown. In eukaryotes, transcription of a given gene usually does not start at a fixed nucleotide position but is distributed across a wider region (33, 34). Genome-wide profiling in humans, mice, and *Arabidopsis* has revealed that dispersed TSSs and alternative TSS usage are widespread, generating transcript isoforms with distinct 5′ ends (33–36). In *Arabidopsis*, alternative TSSs have been shown to exclude uORFs from mRNAs in 220 genes, increasing translation of their mORFs when plants transition from darkness to blue light (35). Similarly, during yeast meiotic differentiation, 380 genes use alternative TSSs to switch between canonical translatable mRNAs and 5′ extended isoforms carrying uORFs that reduce protein synthesis (36). Despite these findings, how the dispersed distribution of TSSs affects uORF regulation under steady-state conditions remains unclear.

We previously identified translated uORFs (TuORFs) in *Arabidopsis* seedlings using improved super-resolution Ribo-seq data with two complementary computational pipelines to detect both long and short TuORFs (24, 37). This comprehensive catalog of 7,051 TuORFs from 4,775 genes enables us to investigate the activity and regulation of diverse TuORFs. Here, we report that most TuORFs in *Arabidopsis* are strongly influenced by transcription start sites (TSSs). This TSS-controlled uORF presence in mRNAs is selective and tightly linked to uORF repressiveness and gene functions. Our findings demonstrate that uORF-mediated translational control is shaped by transcriptional regulation through dispersed TSSs.

## RESULTS

### Analysis of uORF impact reveals that weakly translated uORFs exhibit no mORF repression

Our previous work identified 7,051 TuORFs from 4,775 genes in *Arabidopsis* seedlings (**Table S1A**) (37), enabling us to survey the activity and potential regulation of diverse TuORFs. To quantify TuORF-mediated repression of mORFs, we defined <u>u</u>ORF Impact <u>F</u>actor (uIF) for each TuORF gene based on normalized Ribo-seq ratios between TuORFs and their mORFs (Materials and Methods, **Table S1A**). A higher uIF indicates higher translation in the TuORF relative to the mORF (**Figure 1A**), and *vice versa*. We first focused on 3,213 genes with a single TuORF (**Table S2**) to avoid confounding effects of multiple TuORFs (24). Most TuORFs (62%, 1,995/ 3,213) have a uIF lower than 1 (median = 0.61, **Figure S1**). Consistent with the expectations that more highly translated uORFs exhibit stronger repression, uIF is inversely correlated with mORF translation efficiency (TE) (r = -0.51, **Figure 1B**), supporting uIF as a proxy for TuORF repressiveness. Based on uIF, we grouped the 3,213 TuORF genes into strong (uIF > 1), medium (0.2 <uIF < 1), and weak TuORFs (uIF < 0.2) (**Figure 1C**, **Table S2**). While strong and medium TuORFs are associated with reduced mORF TE, weak TuORFs show no detectable repression compared with genes lacking TuORFs (**Figure 1C**).

**Figure 1:**
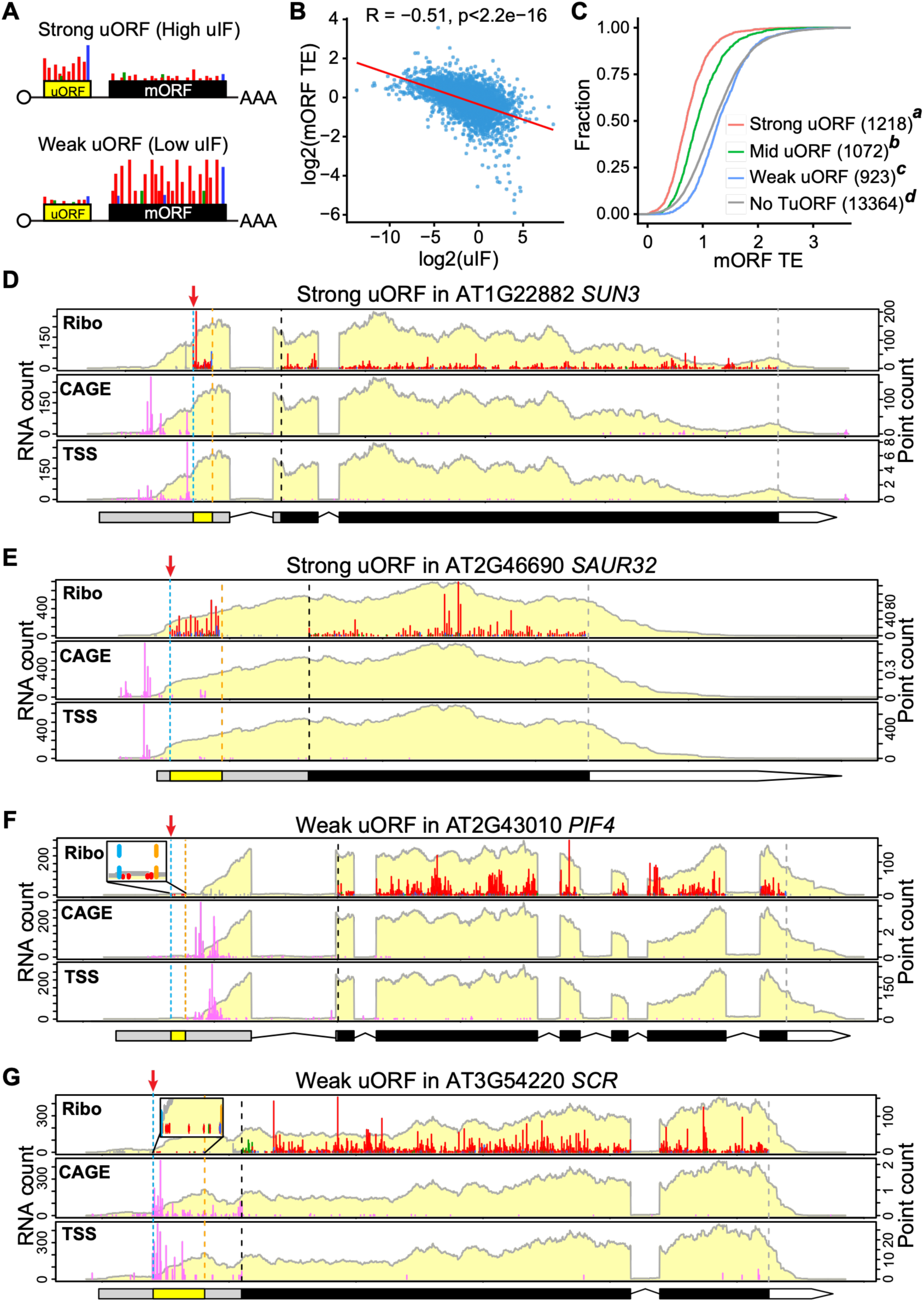
The impact of uORFs is linked to transcription start site positioning. (A) Illustrations of strong and weak uORFs based on uORF impact factor (uIF). (B) Correlation plot of mORF translation efficiency (TE) versus uIF. (C) Cumulative plot of mORF TE in different uIF groups. The “No TuORF” group excluded both RiboTaper- and CiPS-identified TuORF genes. Different italic superscript letters indicate statistical significance between groups (Kolmogorov–Smirnov [KS] test, p < 0.05). (D-G) Multi-omic expression profiles of strong TuORFs (D-E) and weak TuORFs (F-G). Red arrows highlight the uORF start codon; note its position relative to the TSSs detected by CAGE-seq or TSS-seq, between strong and weak uORFs. RNA-seq coverage is shown with a light-yellow background. Ribo-seq, CAGE-seq, and TSS-seq reads are presented by single nucleotide position (point count). Ribo-seq reads are positioned at the first nucleotide of the peptidyl tRNA binding site (P-site) within ribosomes and color-coded as red, blue, or green, to denote the first (expected), second, and third reading frames, respectively; reads falling outside of the ORF ranges are displayed in gray. CAGE-seq and TSS-seq reads are shown in pink. In gene models, black rectangles represent mORFs, yellow rectangles represent uORFs, and gray and white sections represent 5′ UTRs and 3′ UTRs, respectively. Translation start and stop for mORFs are marked by black and gray vertical dashed lines, respectively, and translation start and stop for uORFs are marked by light blue and orange vertical dashed lines, respectively.

To explore mechanisms underlying the low repression of weak TuORFs, we examined RNA-seq and Ribo-seq profiles across different TuORF groups (‘**Ribo**’ panels for **Figure 1D–E** for strong TuORFs, and **Figure 1F–G** for weak TuORFs). Although we confirmed that weak TuORFs are indeed translated, with clear 3-nucleotide periodicity reflecting ribosomes’ codon-by-codon movement, we found that these uORFs are often located near the 5′ edge of RNA-seq coverage or in regions with much lower RNA-seq coverage compared to the rest of the transcript (**Figure 1F–G**). This positional bias suggests that most mRNAs from these genes may use downstream transcription start sites (TSSs), producing transcripts that exclude the uORFs. To test this possibility, we integrated genome-wide TSS profiling from CAGE-seq (Cap Analysis of Gene Expression) and TSS-seq in *Arabidopsis* seedlings (38, 39) with our data. Consistent with our hypothesis, the major TSSs are frequently located downstream of the start codons of weak TuORFs (**Figure 1F-G**, ′CAGE′ and ′TSS′ panels), thereby excluding these TuORFs from the mRNAs. In contrast, the TSSs in strong TuORF genes are positioned upstream of the uORFs, leading to uORF inclusion in the transcripts (**Figure 1D-E**, ′CAGE′ and ′TSS′ panels).

Together, these TSS-uORF positioning biases suggest that TSS selection may control TuORF activity by differentially including uORFs in transcripts.

### Widespread regulation of uORFs by TSS

To determine how widely TSSs affect uORF inclusion, we calculated the percentage of transcripts including the uORF (%uORF+; **Figure 2A**) for each TuORF gene based on TSS data from CAGE-seq. A 90% uORF+ value indicates that 90% of transcripts from a given gene contain the uORF. Strikingly, fewer than 1% (31/ 3,213) of TuORF genes show 100% uORF inclusion (**Figure 2B**). The remaining >99% of TuORF genes are influenced by TSSs to varying degrees (**Figure 2B**). Specifically, 30%, 26%, and 44% of TuORF genes fall into the <30%, 30– 80%, and 80–100% uORF+ categories, respectively, demonstrating that TSSs have a profound effect on uORF presence in mRNAs. In line with the observation that weak TuORFs tend to lie upstream of major TSSs, weak TuORFs are associated with lower %uORF+ (i.e., the uORFs are often excluded from mRNAs; **Figure 2C**, **Table S1A**). In contrast, strong TuORFs (i.e., higher uIF TuORFs as in **Figure 1C**) are preferentially included in transcripts, as indicated by high %uORF+ values (**Figure 2C**).

**Figure 2:**
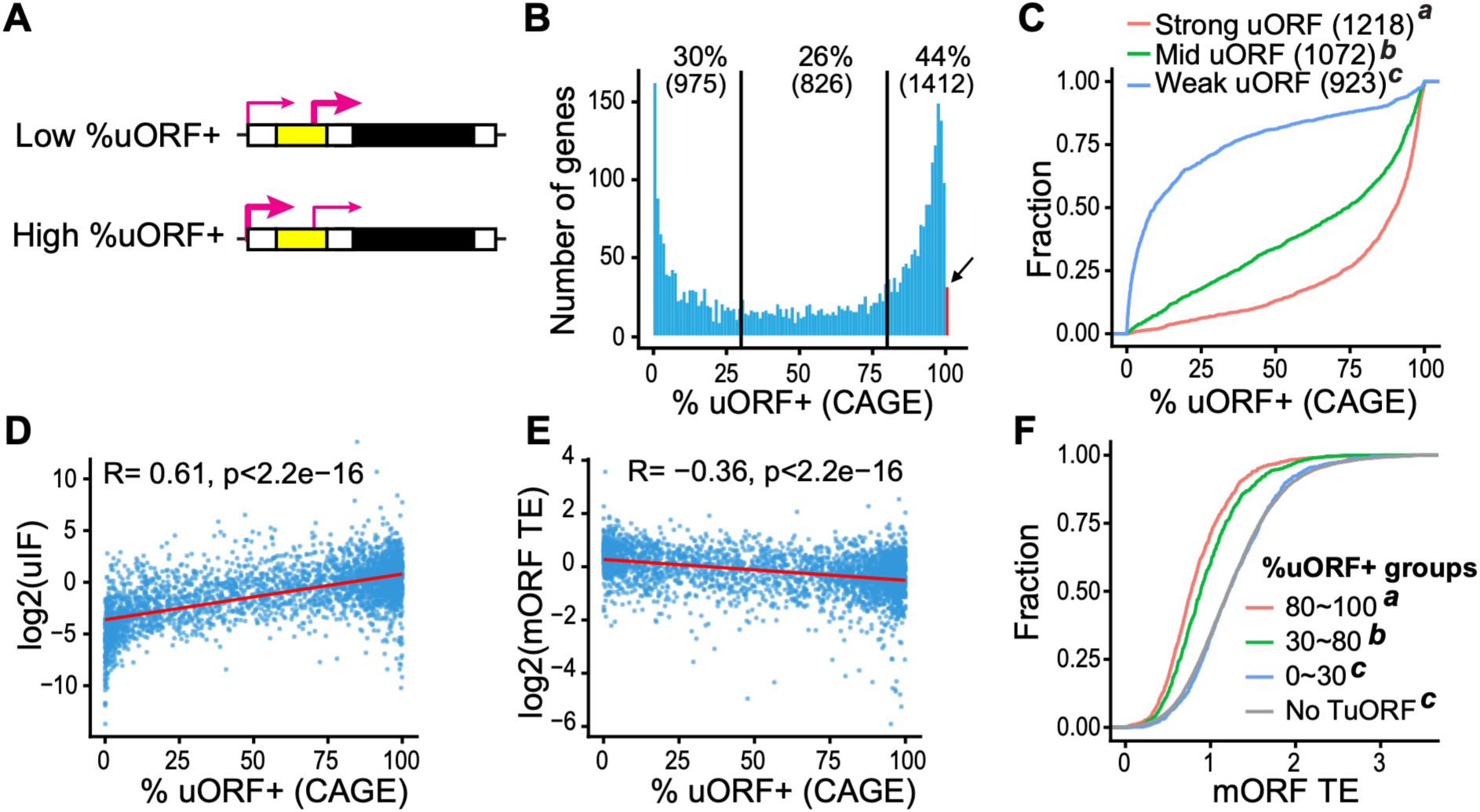
uORF inclusion correlated with uORF repressiveness. (A) The illustration of low and high % uORF inclusion in the transcripts based on TSS choices (curved pink arrows) relative to uORF position. (B) Distribution of % uORF inclusion in each TuORF gene calculated using CAGE-seq data. The red bar highlighted by a black arrow marks the 100% uORF+ genes (<1%). (C) Cumulative plot showing differential uORF inclusion among different uIF uORF groups (as Figure 1C). (D) Correlation plot showing uORF impact is positively correlated with % uORF inclusion. (E) Correlation plot showing mORF TE is negatively correlated with % uORF inclusion. (F) Cumulative plot of mORF TE in different % uORF+ groups. In (C) and (F), different italic superscript letters indicate statistical significance between groups (Kolmogorov–Smirnov [KS] test, p < 0.05).

To assess how TSS positions affect uORF repressiveness, we analyzed the relationships between %uORF+ and uIF or mORF TE. We found that %uORF+ is positively correlated with uIF (r = 0.61, **Figure 2D**) and inversely correlated with mORF TE (r = −0.36, **Figure 2E–F**). These results suggest that higher %uORF+ is associated with stronger uORF- mediated translational repression. Notably, TuORF genes with <30% %uORF+ show no translational repression compared with genes lacking TuORFs (**Figure 2F**). Consistent results were obtained using TSS determined by TSS-seq (**Figure S2A-E**, **Table S1B**). Together, these findings demonstrate that TSS position is a key determinant of uORF inclusion and repressiveness in *Arabidopsis*.

We extended our analysis to genes containing two, three, or four TuORFs. As expected, uORFs located closer to the 5’ end of the transcript (distal to the mORF) are more frequently excluded by downstream transcription start sites (**Figure S3A-C, left panels**); for example, in genes containing two TuORFs, the distal uORF is present in only 61% of transcripts on average, whereas the proximal uORF is present in 91% of transcripts. Consistent with the high exclusion rate, the distal uORFs are associated with lower uIF values (**Figure S3A-C, right panels**). In contrast, mORFs are less affected by TSS usage (**Figure S4**). These results suggest that TSS heterogeneity acts as a progressive filter in complex 5’ leaders, where distal uORFs are frequently excluded while proximal uORFs are more likely to be retained to regulate the mORF.

A related question is why thousands of predicted uORFs have not been detected as translated in deep-coverage Ribo-seq data (24). Are these undetected uORFs (UuORFs) genuinely skipped by ribosomes, or are they excluded from mRNAs due to TSS positions? We examined genes with UuORFs that have appreciable mRNA levels (TPM ≥ 1) (**Table S3**). Like weak TuORFs, but to a greater extent, UuORFs are almost exclusively located upstream of TSSs (**Figure S5A–C**) and are therefore largely excluded from mRNAs. Specifically, 90% of UuORF genes (2,440/2,713) have low %uORF+ (0–30%; **Figure S5D**). The remaining 4.6% (n = 126) and 5.4% (n = 147) of UuORF genes show medium and high %uORF+, respectively (**Figure S5D**), and higher %uORF+ is associated with stronger translational repression (**Figure S5E, left**; **Table S3**). Notably, these uncommon UuORF genes with medium and high %uORF+ generally have low mRNA levels (**Figure S5E, right**), which likely explains why they were not detected in deep Ribo-seq data (24). Together, these results indicate that most UuORF genes (90%) use TSSs downstream of the uORFs, excluding uORFs from mRNAs, while the remaining 10% have low mRNA abundance, limiting their detection.

### TSS positions shape ATG frequency and TuORF inclusion in the transcriptome

Previous studies show that the frequency of ATGs within 5′ UTRs is lower than expected, consistent with purifying selection against uORFs (7, 40–42). However, these analyses were based on annotated gene models or individual cDNAs and did not consider dispersed TSSs.

Because purifying selection against uORFs can act only on transcribed ATGs, ATG depletion should begin at the actual TSSs rather than at annotated TSSs. To validate our finding that many uORFs are excluded from mRNAs by TSSs, we examined ATG frequency around TSSs. We selected the 50% cumulative CAGE-seq positions as a TSS reference point, where the TATA-box and pyrimidine–purine signals corresponding to the transcription initiator motif were observed (**Figure S6**), consistent with previous findings in mammals and *Arabidopsis* (33, 39). We analyzed ATG frequency upstream and downstream of this TSS reference point (**Figure 3A**). In TuORF-less genes, there is a clear transition in ATG frequency across the TSS, with higher frequency upstream (promoters) and lower frequency downstream (5′ UTRs). In TuORF- containing genes, this transition remains significant but is less pronounced (**Figure 3A–B**), in line with the retention of ATGs in 5′ UTRs that give rise to potential uORF(s) and with weaker purifying selection against uORFs in these genes. Together, these results are consistent with TSS-mediated uORF inclusion/exclusion through changes in ATG frequency around TSSs and indicate that TSS positions determine which ATGs are subject to purifying selection against uORFs over evolutionary time.

**Figure 3.**
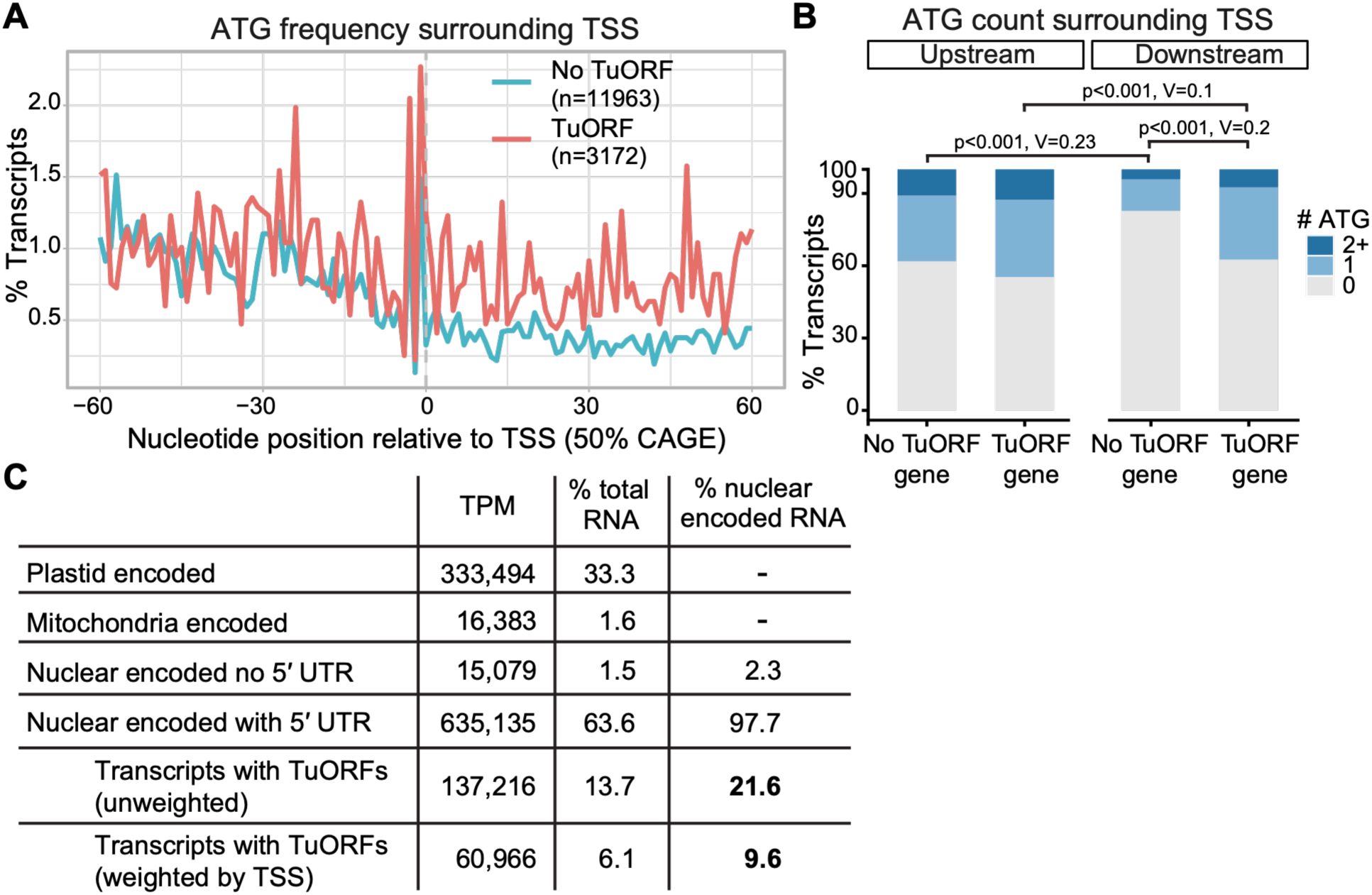
TSS position modulates ATG frequency and affects TuORF inclusion in the transcriptome. (A) ATG frequency surrounding the reference transcription start sites (50% cumulative CAGE position) in No-TuORF and TuORF genes. (B) Proportion of genes with 0, 1, or ≥2 ATGs upstream and downstream of the transcription start sites. Statistical significance was evaluated using a chi-square test comparing the distribution of ATG counts between groups. Effect size was quantified using Cramer’s V (0–1), with ∼0.1, ∼0.3, and ∼0.5 corresponding to small, medium, and large effects. (C) Transcript abundance summary showing that TSS weighting greatly reduces the fraction of nuclear-encoded transcripts that actually contain TuORFs.

### Fewer than 10% of steady-state transcripts contain TuORFs

Although 55% of *Arabidopsis* genes contain potential uORFs (1), our results suggest that the fraction of the transcriptome affected by uORFs is much smaller due to prevalent TSS regulation. We therefore sought to quantify the fraction of transcripts containing TuORFs in the steady-state mRNA pool. For this analysis, we considered all 7,051 TuORFs from 4,775 genes, representing approximately 20% of expressed genes (i.e., 23,862 genes with RNA-seq TPM > 0) (24). When overall mRNA abundance (TPM) is considered, TuORF-containing genes contribute 21.6% of *Arabidopsis* nucleus-encoded mRNAs (**Figure 3C**). However, after accounting for transcripts in which uORFs are excluded by TSS selection (with CAGE-seq data), we found that only 9.6% of nucleus-encoded transcripts contain TuORF(s) (**Figure 3C**). A similar analysis with TSS-seq data identified that only 9.5% of nucleus-encoded transcripts contain TuORF(s) (**Table S4**). These results indicate that, despite the large number of predicted uORF-containing genes and the substantial set of detected TuORFs, only a smaller fraction of the transcriptome is subject to active translational regulation by TuORFs.

### Selective regulation of TuORF genes by TSS

To determine what functions of TuORF genes are differentially regulated by TSSs, we performed Gene Ontology (GO) enrichment analysis on genes containing various %uORF+ categories. We found that different %uORF+ groups enrich functionally distinct gene sets (**Figures S7 to S9**). Genes associated with low uORF+ (< 30%) are significantly enriched for housekeeping functions, including glycolytic enzymes, cell wall components, protein folding, and mitochondrial and chloroplast components. In contrast, genes with high uORF+ (>80%) or multiple TuORFs are preferentially associated with regulatory roles, such as protein kinases, signaling proteins, and transcription factors. The medium uORF+ group (30–80%) displays an intermediate profile, showing enrichment for a mixture of regulatory, chloroplast, and metabolic proteins. These results suggest that the %uORF+ metric can be a predictor of a transcript’s functional identity, distinguishing stable housekeeping genes from regulatory genes.

We next asked whether TSSs are adapted to beneficial uORFs, such as evolutionarily conserved CPuORFs, to ensure their inclusion in mRNAs to carry out their regulatory functions. We analyzed the 67 CPuORF genes that were expressed in our data. We found that CPuORFs are predominantly located downstream of the TSSs defined by CAGE-seq and TSS-seq (**Figure 4A–C**), with most CPuORF genes exhibiting ≥ 90 % uORF+ (**Figure 4D**). Compared to other TuORFs, CPuORFs exhibit a significantly higher uIF (**Figure 4E**) and show stronger repression of their mORFs (**Figure 4F**). These observations suggest that TSSs may have coevolved with CPuORFs (and other strong uORFs, **Figure 2C**) to preserve regulatory uORFs in transcripts.

**Figure 4:**
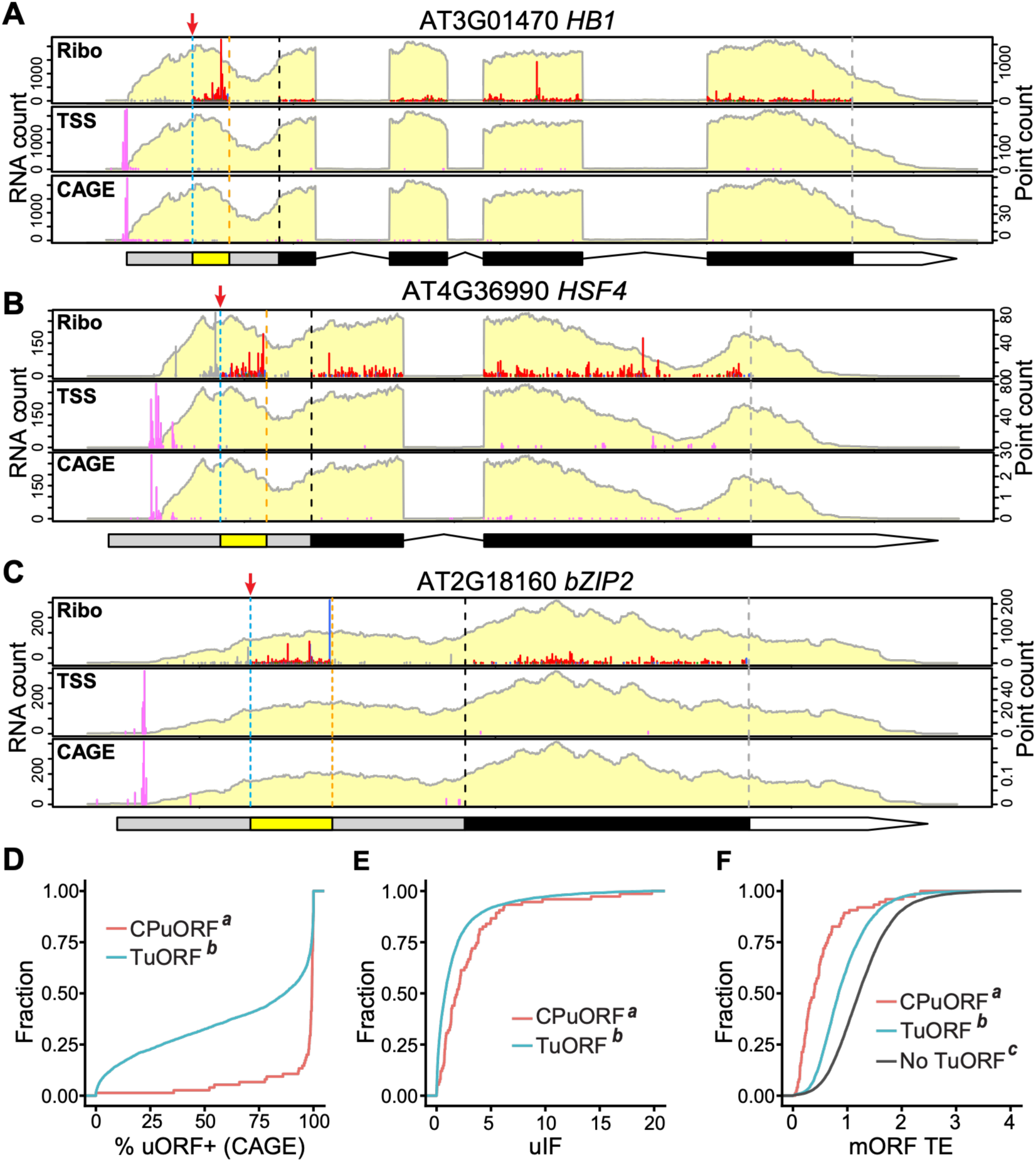
Conserved peptide uORFs (CPuORFs) are preferentially preserved within the mRNAs. (A–C) Expression profiles of example CPuORF genes. The data are presented as described in the Figure 1 legend. (D) Cumulative plot of %uORF inclusion in CPuORF genes and other TuORF genes. (E) Cumulative plot of uIF for CPuORF genes and other TuORF genes. (F) Cumulative plot of mORF TE comparing CPuORF genes, other TuORF genes, and genes without TuORFs. In (D–F), different italic superscript letters indicate statistical significance between groups (Kolmogorov–Smirnov [KS] test, p < 0.05).

Together, these findings suggest that TSSs selectively regulate the inclusion of TuORFs in mRNAs, thereby tailoring uORF-mediated translational control to genes with distinct biological functions.

### Long uORFs show stronger repression after accounting for TSSs

Given that ∼1/3 of TuORF genes with ≤ 30% uORF+ show no translational repression (**Figure 2F**, **Figure S2E**), which may confound interpretation of uORF activity, we revisited how uORF repressiveness is controlled by uORF length. Because longer uORFs can impede ribosome reinitiation, longer uORFs are expected to exhibit stronger repression than shorter ones (43, 4). However, this length-dependent repression is obscured when considering all TuORFs (**Figure 5A**) (24). By focusing on TuORFs that are included in greater than 30% of the transcripts, a clear length-dependent uORF repression pattern was observed, and longer uORFs exhibit stronger repression of mORFs (**Figure 5B**). Notably, the 1-aa uORFs deviate from this length-dependent trend (**Figure 5B**), possibly because they often induce strong stalling effects and block translation (44, 45).

**Figure 5:**
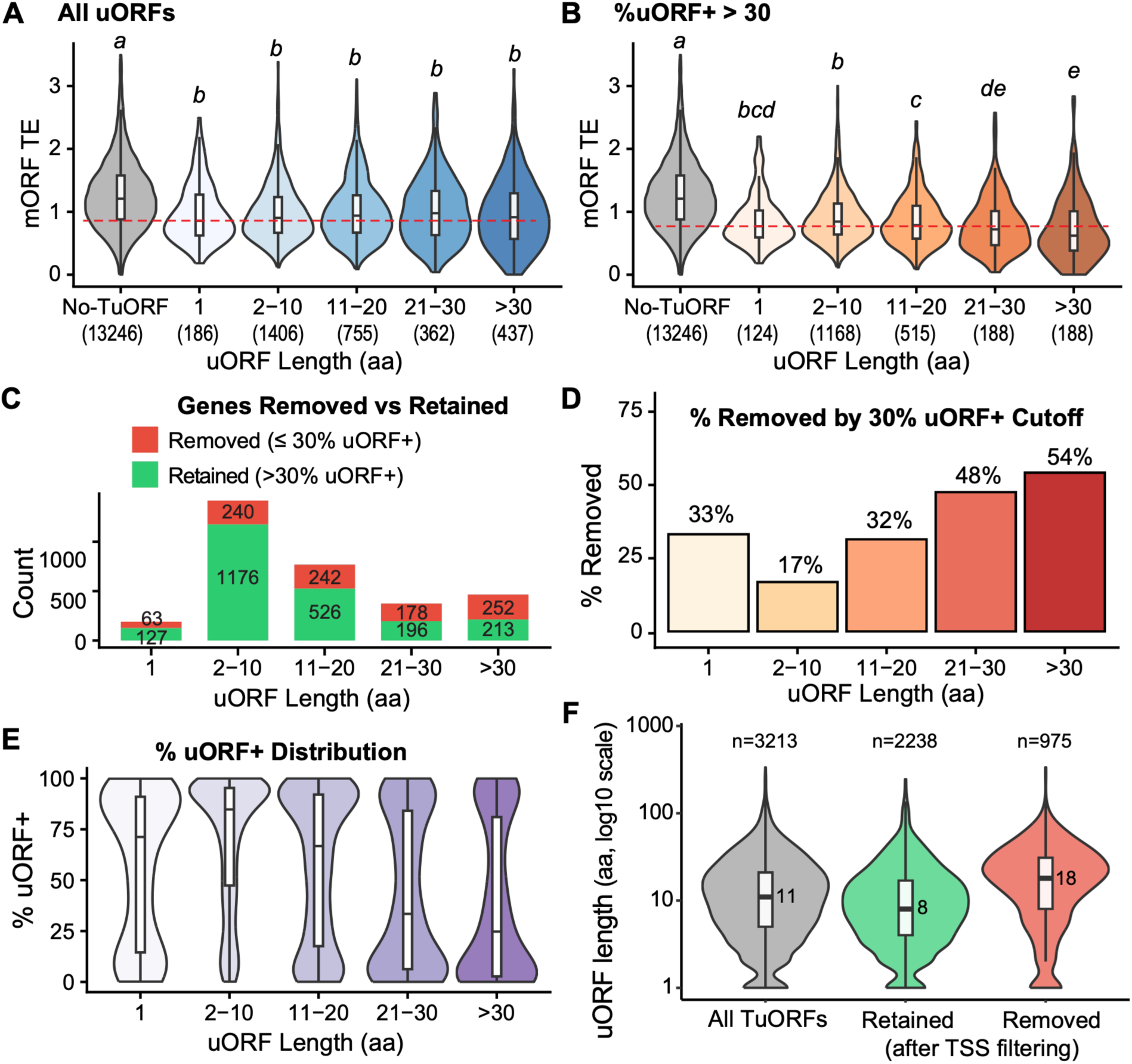
Accounting for TSS reveals that longer uORFs are more repressive than short uORFs. (A–B) mORF TE of genes containing different lengths of TuORFs without considering TSSs (A) and after filtering TuORF genes ≤ 30% uORF inclusion (B). The red dashed line marks the median of the 1-aa uORF group. (C) Number of genes removed (≤30% uORF+) or retained (>30% uORF+) in each uORF length group. (D) Percentage of genes removed by the 30% uORF+ cutoff in each uORF length group. (E) Distribution of %uORF+ across uORF length groups, illustrating progressively lower uORF inclusion for longer uORFs. (F) uORF length distributions for all TuORFs and for TuORFs retained or removed after TSS filtering. Numbers beside the boxes indicate median uORF length (aa).

We found that the proportion of uORFs excluded by TSSs (≤30% uORF+) increases progressively with uORF length (**Figure 5C, D**). While short uORFs are mostly preserved in transcripts (17% excluded for uORF 2–10 aa), the uORF exclusion rate rises sharply for longer uORFs (32%, 48% and 54% excluded for uORFs 11–20 aa, 21–30 aa and > 30 aa) (**Figure 5C, D**). Again, the 1-aa uORFs (n = 190; 33% excluded) deviate from the length-dependent trend, possibly due to their strong repressive effects (**Figure 5C, D**). The distribution of %uORF+ transcripts shifts from high inclusion for short uORFs to broad exclusion for longer ones (**Figure 5E**). As a result, after TSS filtering, the overall TuORF length is reduced (**Figure 5F**). It is possible that longer uORFs are more likely excluded by chance due to their length, or they are excluded due to higher purifying selection against stronger repressors.

Together, these findings demonstrate that accounting for TSS usage is essential for revealing the expected length-dependent repression by uORFs and that TSS heterogeneity preferentially excludes long, strongly repressive uORFs.

### Weak uORFs and undetected uORFs can suppress translation in artificial conditions, leading to misinterpretation of uORF activity

Although weak and undetected uORFs (UuORFs) are largely excluded from mRNAs in vivo, we investigated whether they exhibit repressive activity when artificially included in the 5’ UTR. We characterized four weak uORFs from *SVBL*, *PIF4*, *PAPS1*, and *IAR1*, and two UuORFs from *UL14X* and *NPQ4* using a dual-luciferase assay. We found that all these uORFs, except the one from *PAPS1*, significantly repress downstream *FLUC* translation compared with mutated uORFs (start codon ATG>**TA**G) (**Figure 6A**). The uORF from *PAPS1* slightly reduces FLUC, but the effect was not statistically significant (**Figure 6A**). We reasoned that the weaker repression by the *PAPS1* uORF may result from its poor Kozak context (ACGA<u>ATG</u>TT) compared with the optimal consensus sequence (AAAA<u>ATG</u>GC) (46). When we modified the uORF Kozak sequence to the optimal (strong) context for *PAPS1*, *SVBL*, and *IAR1*, these uORFs exhibited increased repressive activity (**Figure 6B**). Together, these results demonstrate that artificially introducing a uORF that is typically absent from transcripts can still suppress downstream translation. These findings highlight that such uORFs may be misinterpreted as biologically functional when TSS is not considered.

**Figure 6:**
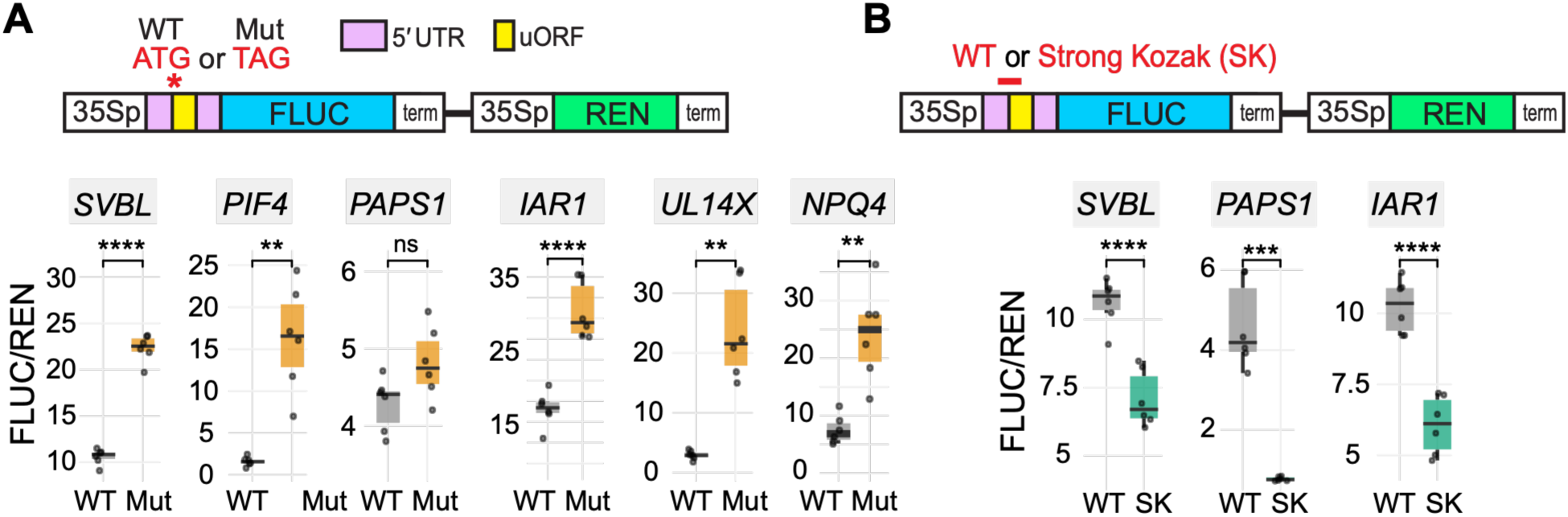
uORFs that are normally absent from mRNAs still repress downstream translation when artificially included. (A) Dual luciferase assay comparing wildtype and mutated versions of weak uORFs in *SVBL*, *PIF4*, *PAPS1*, and *IAR1*, and untsaranslated uORF in *UL14X* and *NPQ4*. The annotated 5′ UTR for each gene carrying the wild-type uORF or mutated uORF (start codon ATG>**TA**G) was synthesized and cloned upstream of the *FLUC* reporter. FLUC luminescence levels were normalized to REN luminescence levels. (B) Dual luciferase assay examining whether a strong Kozak sequence further improves uORF strength for weak uORFs in *SVBL*, *PAPS1*, and *IAR1*. The statistical significance was determined by two-sample Student’s t-test (n=6, * p<0.05, ** p<0.01, *** p<0.001, **** p<0.0001).

## DISCUSSION

Our study identifies TSS heterogeneity as a major determinant of uORF activity in *Arabidopsis* and reveals that translational control by uORFs is shaped by transcriptional regulation through TSS choices. Our findings also help explain the longstanding paradox of why uORFs remain prevalent at the genome sequence level despite their potent repressive potential. We showed that *Arabidopsis* uses dispersed TSSs to generate transcripts that often exclude uORFs from 5′ UTRs, thereby reducing the proportion of transcripts that are subject to uORF- mediated repression (**Figure 7**). This TSS-mediated filtering for uORF inclusion is selective and linked to gene functions. uORFs encoding conserved peptides or associated with genes encoding regulatory proteins tend to be retained in mRNAs, whereas uORFs associated with genes encoding housekeeping proteins are more likely to be excluded. Importantly, after removing uORFs that are rarely present on mRNAs from the analysis, the expected repressive effect of uORF length becomes clear, revealing that longer retained uORFs are associated with stronger inhibition of mORFs. Longer uORFs are also more frequently excluded, consistent with selection against extreme translational inhibition. In genes with multiple uORFs, TSS heterogeneity further creates a positional hierarchy in which distal uORFs are more often excluded. Consistently, the relative position of a uORF to the TSS in the 5′ UTRs could significantly affect its activity (47). Together, these findings support that TSS heterogeneity is an important regulatory layer that determines uORF activity. Since TSSs are known to be regulated under different growth conditions and developmental stages in diverse organisms (34–36), conditional uORFs could contribute to the reprogramming of gene expression through TSS changes during developmental processes and environmental responses. Because dispersed TSS usage is a general feature of eukaryotic promoters, TSS-mediated filtering of uORFs is likely to operate well beyond *Arabidopsis*.

**Figure 7.**
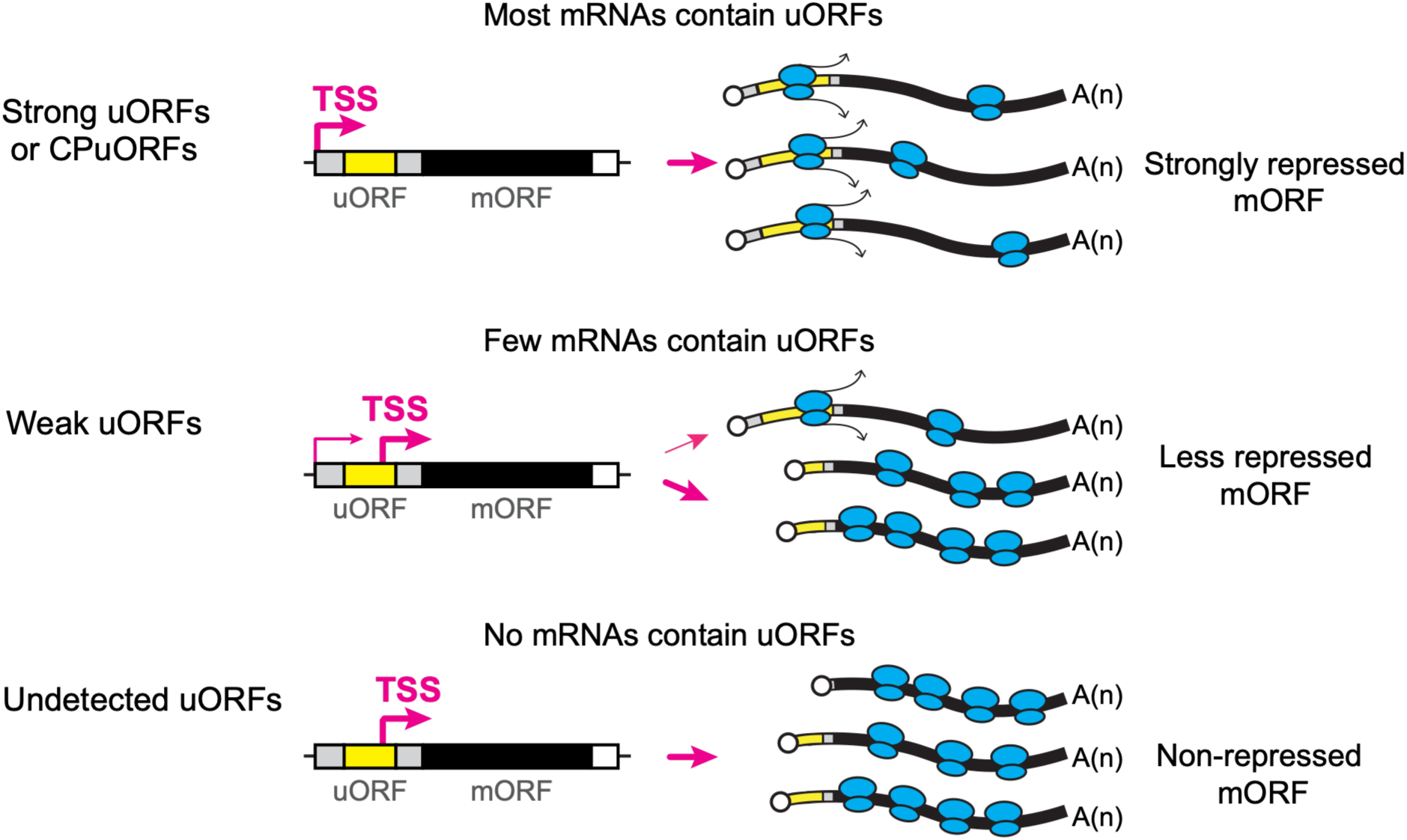
Model for how dispersed TSSs affect uORF repressiveness. A model for TSS regulation of uORF repressiveness. The curved pink arrows indicate the TSSs. In strong uORF or CPuORF genes, upstream TSSs are preferred, allowing high uORF inclusion in the mRNAs, resulting in strong repression toward mORFs. In contrast, in weak uORF genes, downstream TSS is favored over upstream TSS, producing transcripts mostly lacking uORFs, resulting in weak repression of mORFs overall. Most undetected uORFs are exclusively excluded from mRNA sequences and do not repress their mORFs.

In biology, we often simplify our thinking by focusing on the number of genes involved in a process without considering their transcript abundance. Yet, the number of genes could be disproportionate to their transcript abundance. For example, while only 5% of yeast genes contain introns, these genes produce up to 27% of total transcripts in the genome (48). Although 55% of *Arabidopsis* genes contain potential uORFs based on the annotated 5′ UTRs, after considering the fraction of uORFs eliminated by TSSs and transcript abundance, only 9.6% of the mRNA pool harbor potential TuORFs. Note that uORFs that are too close to the TSSs may not initiate translation efficiently due to poor scanning (49, 50). Hence, the actual fraction of *Arabidopsis* mRNAs regulated by uORFs is likely even lower than 9.6%.

In addition to uORFs, our work provides a warning note for studying other mRNA features in 5′ UTRs, such as secondary structures or binding motifs for RNA-binding proteins (51). Because modern genome annotations incorporate deep RNA-seq data that capture rare 5′ extensions, the annotated 5′ UTRs are often longer than those predominantly used by abundant mRNA isoforms in vivo. Analyses based solely on annotated 5′ UTRs can lead to misidentification and misinterpretation of features that are generally absent from the actual transcriptome. At least a fraction of undetected uORFs in *Arabidopsis* could result from the Araport11 annotation overestimating 5′ UTR lengths, as previously reported (39, 52). We demonstrated that artificially expressed uORFs that are generally absent from mRNAs can still repress downstream translation, which can lead to misinterpretation of uORF activity in vivo. Therefore, we recommend that researchers studying 5′ UTR features incorporate TSS heterogeneity into their analyses. When TSS resources are unavailable, cross-referencing RNA-seq coverage with annotated gene models helps detect if the annotated 5′ UTR is overextended. In addition, filtering out uORFs and other mRNA features with low functional impact, as indicated by low uIF values for uORFs, can improve the identification and characterization of biologically relevant regulatory elements. Finally, beyond routine quantification of expression levels, we urge researchers to visualize TSS distributions alongside gene expression profiles (53) to integrate transcriptional and post-transcriptional regulation and gain a more comprehensive and accurate understanding of gene regulation.

## MATERIALS AND METHODS

### Data source of TuORFs and software used in this study

The translated uORFs (TuORFs) were extracted from our previous high-coverage super-resolution Ribo-seq data in 7-day-old *Arabidopsis* seedlings (24). 2,106 TuORFs were uniquely identified by RiboTaper (54), and 6,627 TuORFs were uniquely identified by CiPS; 1,682 TuORFs were identified by both methods, resulting in a total of 7,051 TuORFs (**Table S1A**). The TuORFs identified here are slightly higher than those previously identified by CiPS (24) because we now allow CiPS to include TuORFs whose stop codon overlaps the mORF start codon, to be consistent with RiboTaper.

Data analysis was accomplished with R 4.3.0 (55). Plots for RNA-seq, Ribo-seq, CAGE-seq, and TSS-seq were visualized by *RiboPlotR* (56). Other plots were created with *ggplot2* (57).

### Quantification of <u>u</u>ORF Impact <u>F</u>actor (uIF)

We extracted the Ribo-seq P-site counts of the TuORFs and their corresponding mORFs. Next, we normalized the read counts with the ORF length. <u>u</u>ORF Impact <u>F</u>actor (uIF) is calculated by the ratio of uORF and mORF normalized Ribo-seq reads. Similar normalization strategies have been applied to study uORF-mediated translational repression (58, 59). For simplicity, only genes containing one TuORF were included in this analysis to avoid cumulative translational repression from multiple uORFs.

### Quantification of CAGE-seq and TSS-seq reads for TuORF genes

CAGE-seq and TSS-seq data from *Arabidopsis* seedlings (38, 39) were extracted from the JBrowse hosted by TAIR (www.Arabidopsis.org) (60). To visualize the TSS and our TuORF expression profiles in parallel, we organized the CAGE-seq and TSS-seq data into tabular files as input for *RiboPlotR* (56), containing chromosome number, strand, position, and read counts. We then visualized RNA-seq, Ribo-seq, CAGE-seq, and TSS-seq together with the ′PLOTch3′ function in *RiboPlotR*. We encourage interested users to visualize omics data using the updated, more user-friendly ggRibo() function in the *ggRibo* package (53).

To calculate the percentage of upstream transcription start sites relative to a given TuORF (%uORF+), we first mapped CAGE-seq or TSS-seq reads to annotated 5′ UTRs with an additional 100 nucleotides upstream. Next, we calculated the total CAGE-seq and TSS-seq counts and the counts upstream of the TuORF to determine the %uORF+.

### Defining undetected uORFs

Genes that contain TuORFs identified either by RiboTaper or CiPS were first removed. The remaining genes with RNA-seq TPM ≥ 1 and containing one potential AUG-initiated uORF≥ 10 codons (including the stop codon) were used for downstream analysis.

### Determining mRNA expression levels and mORF translation efficiency

TPM (transcripts per million) of RNA-seq and Ribo-seq levels were quantified by RSEM (v1.3.1) (61) as described previously (24). The mORF translation efficiency was calculated by dividing Ribo-seq TPM by RNA-seq TPM corresponding to the mORF coding regions.

### Estimating the fraction of transcripts regulated by at least one TuORF

All TuORFs identified by either RiboTaper or CiPS (24) were used for this analysis. For each TuORF gene, we determined the TSS counts within 5′ UTR plus an additional 100 nucleotides upstream. Next, we calculated the %uORF+ for each TuORF gene as follows:

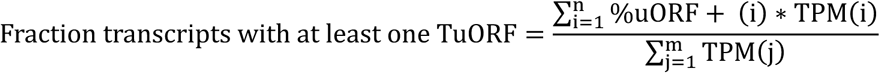

n is the total number of genes containing TuORFs. m is the total number of expressed genes. For genes containing ≥ 2 TuORFs, only the TuORF closest to the mORF was used for calculating this value.

### Identifying CAGE-seq-defined TSSs and calculating AUG frequency surrounding TSSs

For a given gene, we first determined total CAGE-seq reads within the annotated 5′ UTR plus the potential promoter region (i.e., an additional 100 nucleotides upstream). Next, we calculated the cumulative CAGE-seq counts. The CAGE-seq-defined TSS for each gene was defined as the nucleotide position where the cumulative CAGE-seq counts reached 50% of the total CAGE-seq counts. Based on the CAGE-seq-defined TSSs, we calculated the AUG frequency across 120 nucleotides upstream and downstream of the TSSs.

### Dual-luciferase assay

To test the weak and undetected uORFs on downstream translation when forcibly included in the 5′ UTRs, a DNA fragment containing the 35S promoter and 5′ UTR (with the intact uORF or the start codon mutated to TAG) was synthesized and inserted upstream of the *FLUC* reporter in pGreen II 0800-Luc (62) via BamH I and Nco I. For the optimized (strong) Kozak version, the synthesized sequence surrounding the start codon was changed to AAAA<u>ATG</u>GC (46). The exact sequences synthesized were described in **Supplemental File 1.**

The gene synthesis and cloning were performed by BioBasic.

*Arabidopsis thaliana* T87 cell line (rpc00008) was provided by the RIKEN BRC through the National BioResource Project of the MEXT, Japan. The cell lines were maintained in 50 ml liquid media (4.73 g/L LS (Caisson, LSP03-50LT), 3% sucrose, 0.00002% Thiamine (B1), 0.002% Myoinositol, 0.018% KH_2_PO_4_, and 0.2mg/L 2,4-D) in a 250 ml baffled Erlenmeyer flask shaken at 140 rpm at 22°C in the dark and subcultured every 7 days.

The dual-luciferase assay was performed as described in (24) with minor modifications. Briefly, freshly subcultured *Arabidopsis* T87 cells (3 ml 7-day-old stock transferred to 50 ml media) after 3 days were used. The cells were pelleted by centrifugation at 100 g for 3 minutes and resuspended in the enzyme digestion solution for 4 hours to generate protoplasts. The protoplast solution was filtered through a 40 µm cell strainer prior to washing with W5 buffer. 5 µg of the plasmid DNA isolated with the ZymoPURE II Plasmid Midiprep kit (Zymo D4201) was transformed into 10^5 protoplasts using the PEG method. Subsequently, the protoplasts were transferred to 24-well culture plates and incubated overnight in the dark at 22°C. Each plasmid was transformed in 6 biological replicates, and two to three independent experiments were performed. The protoplasts were lysed using Passive Lysis Buffer included in the Dual-luciferase assay kit (Promega E1960), and FLUC and REN luminescence was measured using 20 µL of the cleared lysate with GloMax Navigator Plate Reader (Promega, GM2010). The FLUC luminescence was normalized to its corresponding REN luminescence.

### AI usage

The authors used large language models (Claude Opus 4.7-5.0 Max, Grok 4.1-4.4 Expert, Gemini 3.1 Pro, ChatGPT 4.5 Pro-5.6 Sol Ultra) to assist with manuscript editing for grammar and clarity and to improve analysis code according to authors’ instructions. These tools were not used to generate data or draw scientific conclusions. All outputs were reviewed and verified by the authors, who take full responsibility for the content of the manuscript.

## AUTHOR CONTRIBUTIONS

HLW and PYH designed the research and interpreted the data. HLW analyzed the sequencing data and prepared figures and tables for the manuscript. QA performed the DLA. HLW and PYH wrote the manuscript.

## ACKNOWLEDGEMENTS

We thank Christoph Benning and Linda Danhof for sharing the *Arabidopsis* T87 culture cells and relevant protocols. We thank Isaiah Kaufman for his critical review of this manuscript. This work was supported in part through computational resources and services provided by the Institute for Cyber-Enabled Research at Michigan State University. This work was supported by the National Science Foundation (NSF) under award numbers 2425390 and 2051885 and the National Institute of General Medical Sciences of the NIH under award number R35GM155375 to PYH. The content is solely the responsibility of the authors and does not necessarily represent the official views of the NSF or NIH.

---

**Figure S1:**
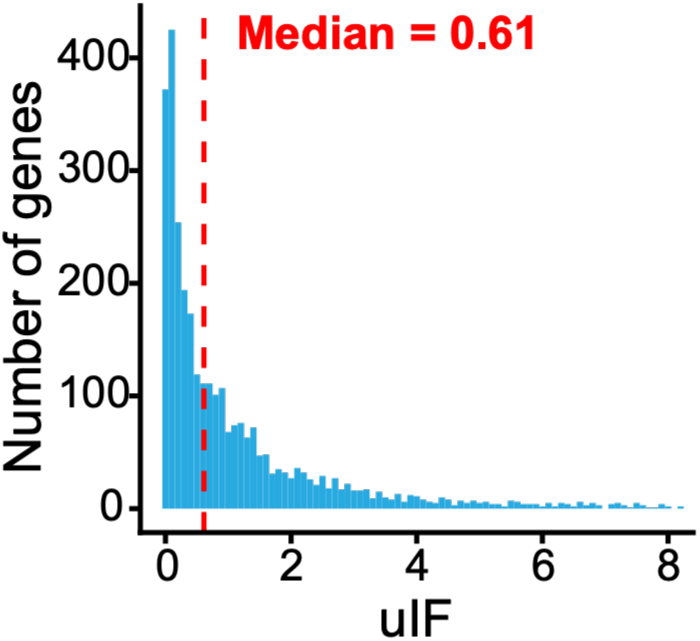
Distribution of uIF among TuORF genes. The vertical dashed line marks the median.

**Figure S2:**
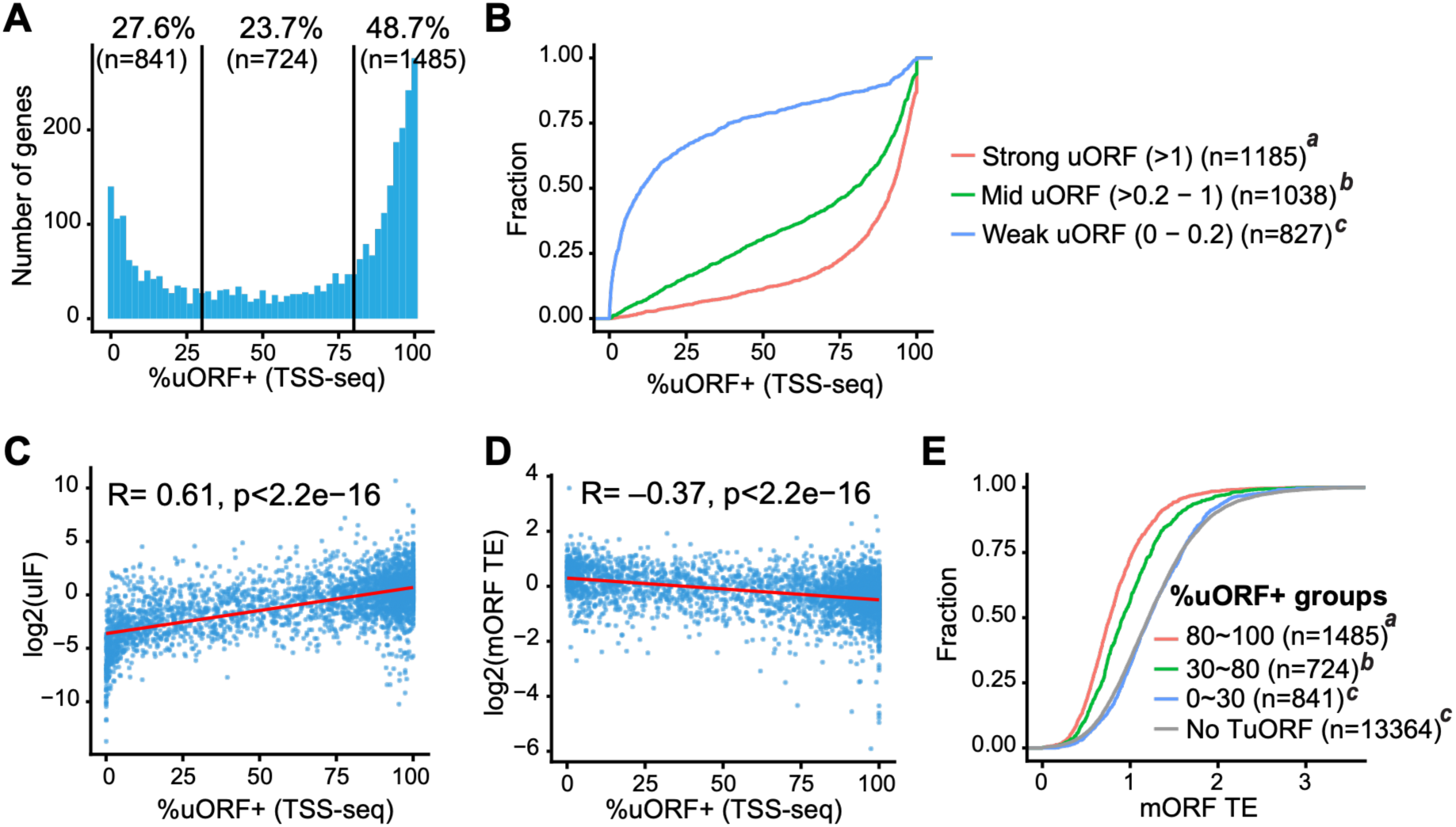
uORF inclusion correlated with uORF repressiveness, shown with TSS-seq data. The same analysis as Figure 2, but using TSS-seq data. (A) Distribution of % uORF inclusion in each TuORF gene calculated using TSS-seq data. Approximately 7% of TuORF genes show 100% uORF inclusion. (B) Cumulative plot showing differential uORF inclusion among uIF uORF groups. (C) Correlation plot showing uORF impact is positively correlated with % uORF inclusion. (D) Correlation plot showing mORF TE is negatively correlated with % uORF inclusion. (E) Cumulative plot of mORF TE in different % uORF+ groups. The “No TuORF” group excluded both RiboTaper- and CiPS-identified TuORF genes. In (B) and (E), different italic superscript letters indicate statistical significance between groups (Kolmogorov–Smirnov [KS] test, p < 0.05).

**Figure S3.**
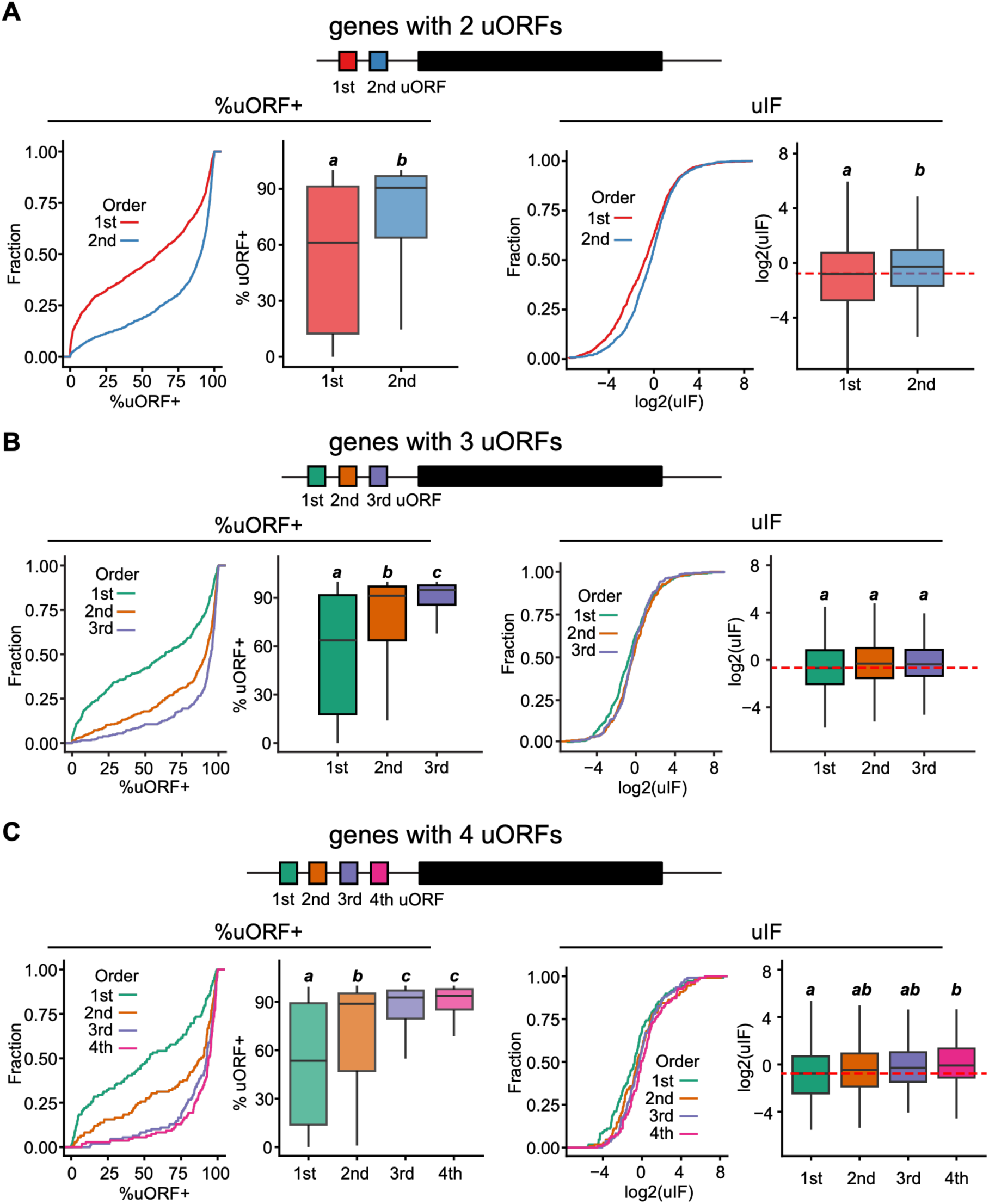
Distal uORFs are preferentially excluded from mRNA in multiple uORF genes. (A–C) Analysis of genes containing 2 uORFs (A, n = 1020), 3 uORFs (B, n = 320), or 4 uORFs (C, n = 109). In each gene, uORFs were ordered from the 5′ end (1^st^: most distal from the mORF) to the mORF-proximal uORF. Cumulative plots and boxplots are shown for %uORF+ (left panels), which measures the proportion of transcripts containing each uORF, and for uIF (right panels), which shows the relative translation levels of each uORF normalized to its mORF. The red dashed lines mark the median of the 1^st^ uORF.

**Figure S4.**
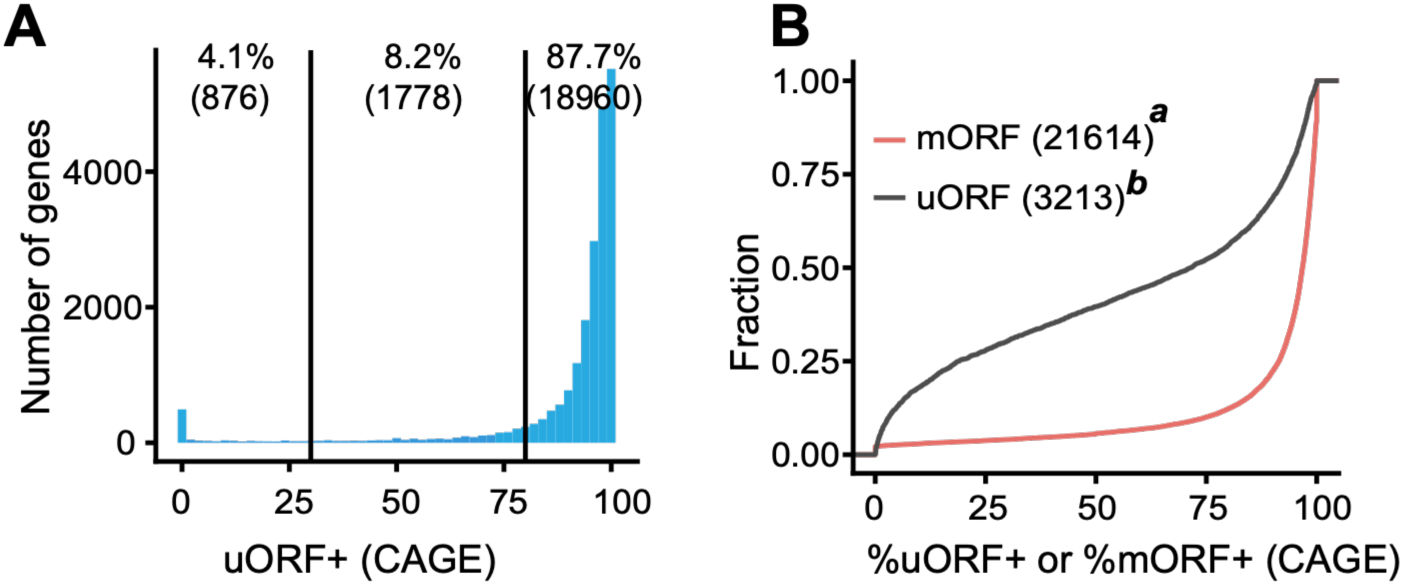
mORFs are less affected by TSS heterogeneity. (A) Distribution of %mORF+ (similar to %uORF+ but calculates the percent of TSS upstream of the mORF) based on CAGE-seq data. (B) Comparison of %uORF+ and %mORF+ based on CAGE-seq data.

**Figure S5:**
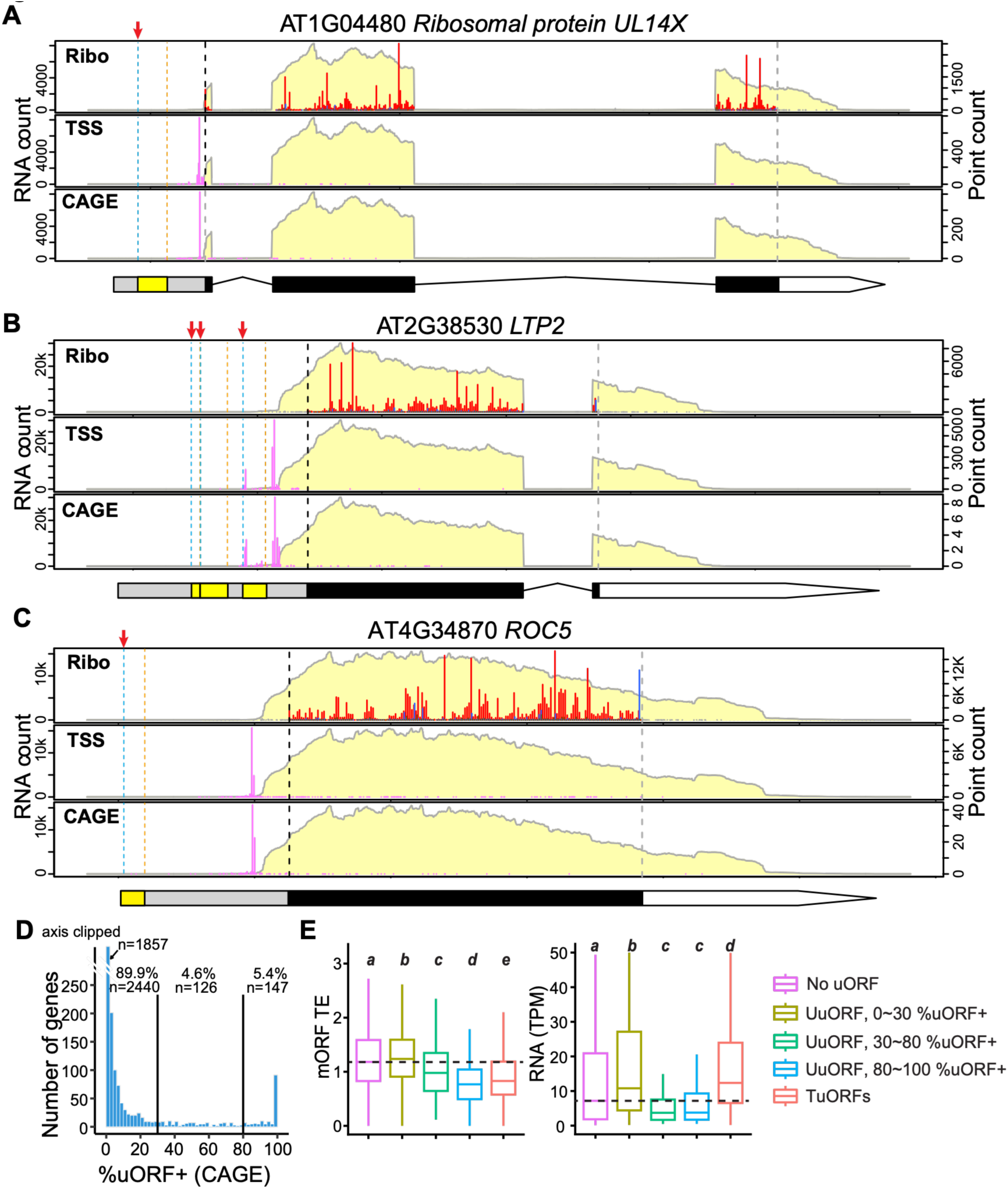
Undetected uORFs are generally excluded from the mRNA sequences. (A–C) Examples of predicted but undetected uORFs (UuORFs). The data are presented as described in the Figure 1 legend. Red arrows highlight the uORF start codon. (D) Distribution of %uORF+ of UuORF genes based on CAGE-seq data. Note that, for UuORF genes, we only considered transcripts containing a single predicted uORF greater than 10 codons in length and undetected in (24). ‘n’ indicates the number of uORF genes in each group. The %uORF+ is based on CAGE-seq data. (E) mORF TE and mRNA levels comparing no-uORF, UuORF genes with varying %uORF+, and TuORF genes. Dashed lines mark the median of the ‘No uORF’ group. In (D-E), different italic superscript letters indicate statistical significance between groups (Pairwise Wilcoxon rank-sum test with Benjamini–Yekutieli correction for FDR).

**Figure S6:**
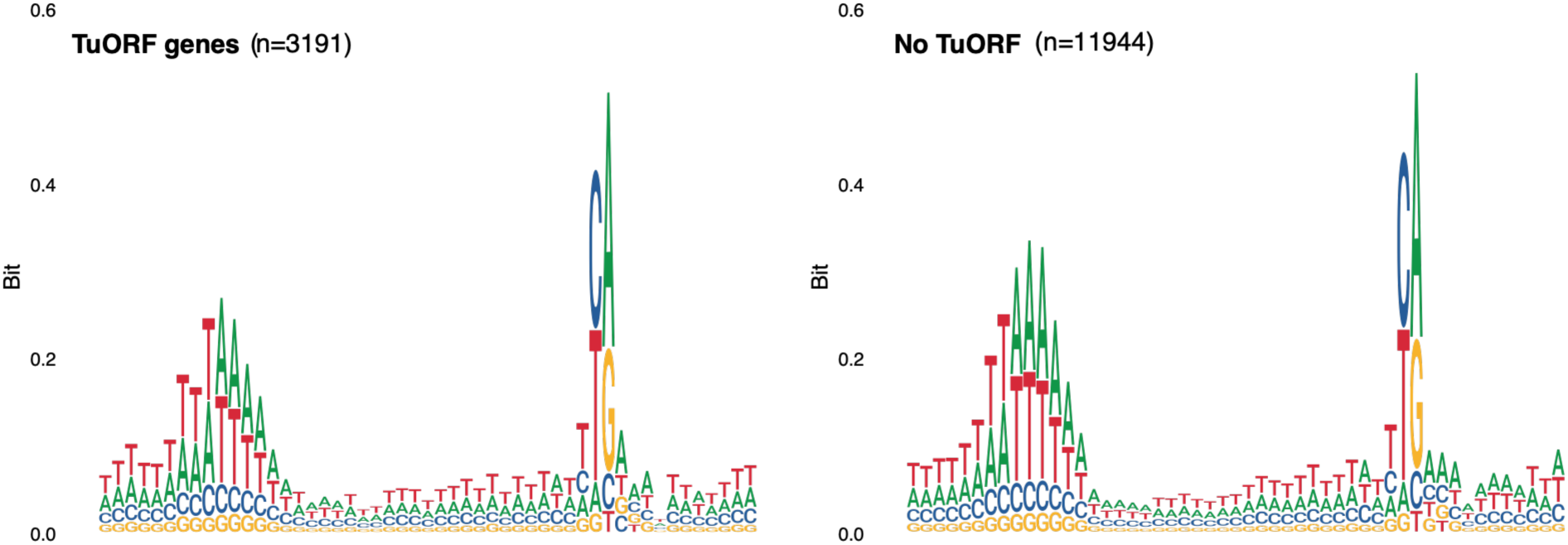
Sequence context of transcription start sites in TuORF and no-uORF genes. Sequence logos showing nucleotide frequencies surrounding the 50% CAGE site in TuORF genes (left) and genes without TuORFs (right). The TSS is centered at position +1, and the sequence logos span 120 nucleotides. Letter heights indicate information content.

**Figure S7:**
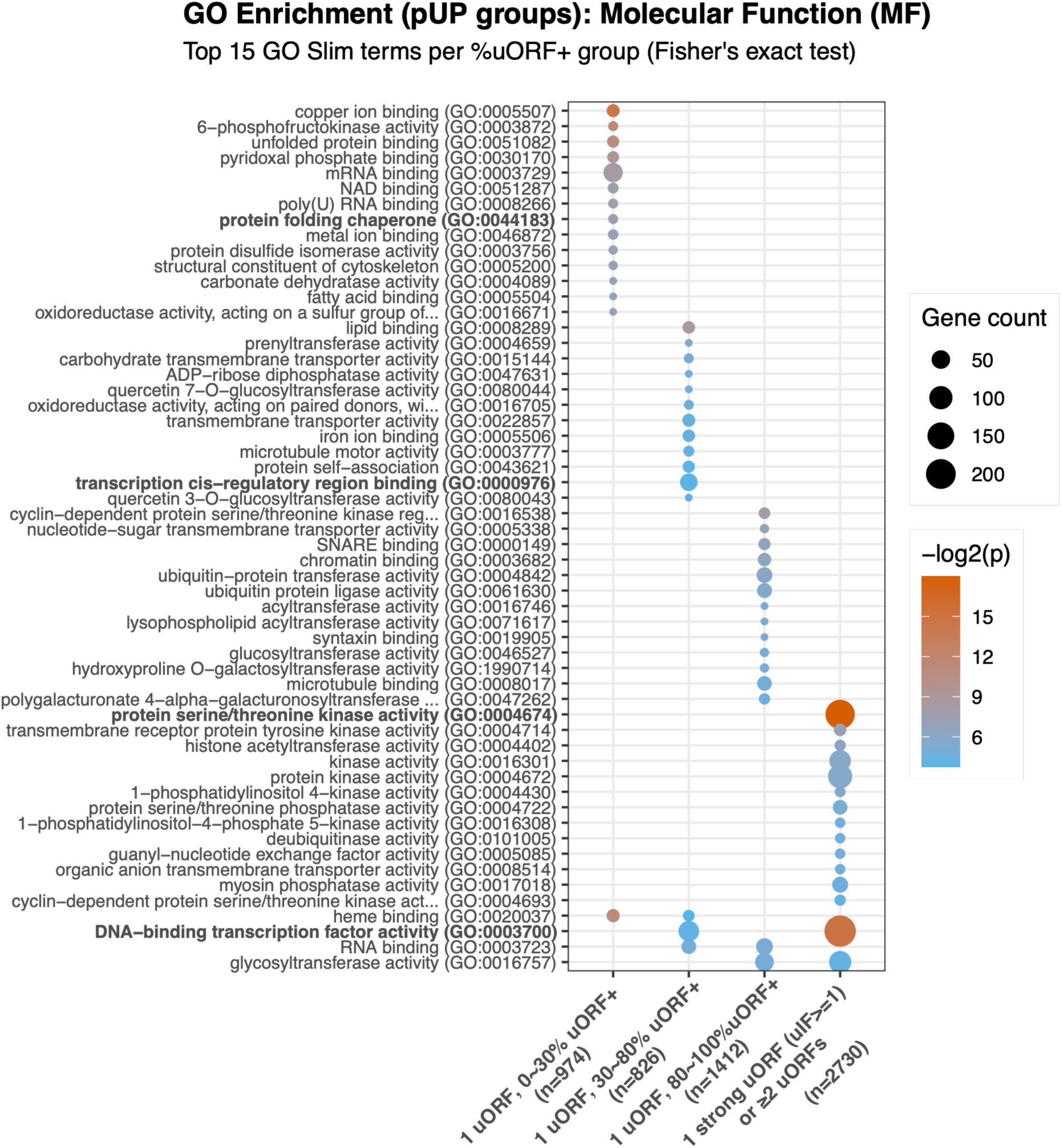
GO Molecular Function terms enriched in uORF groups across different %uORF+ categories.

**Figure S8:**
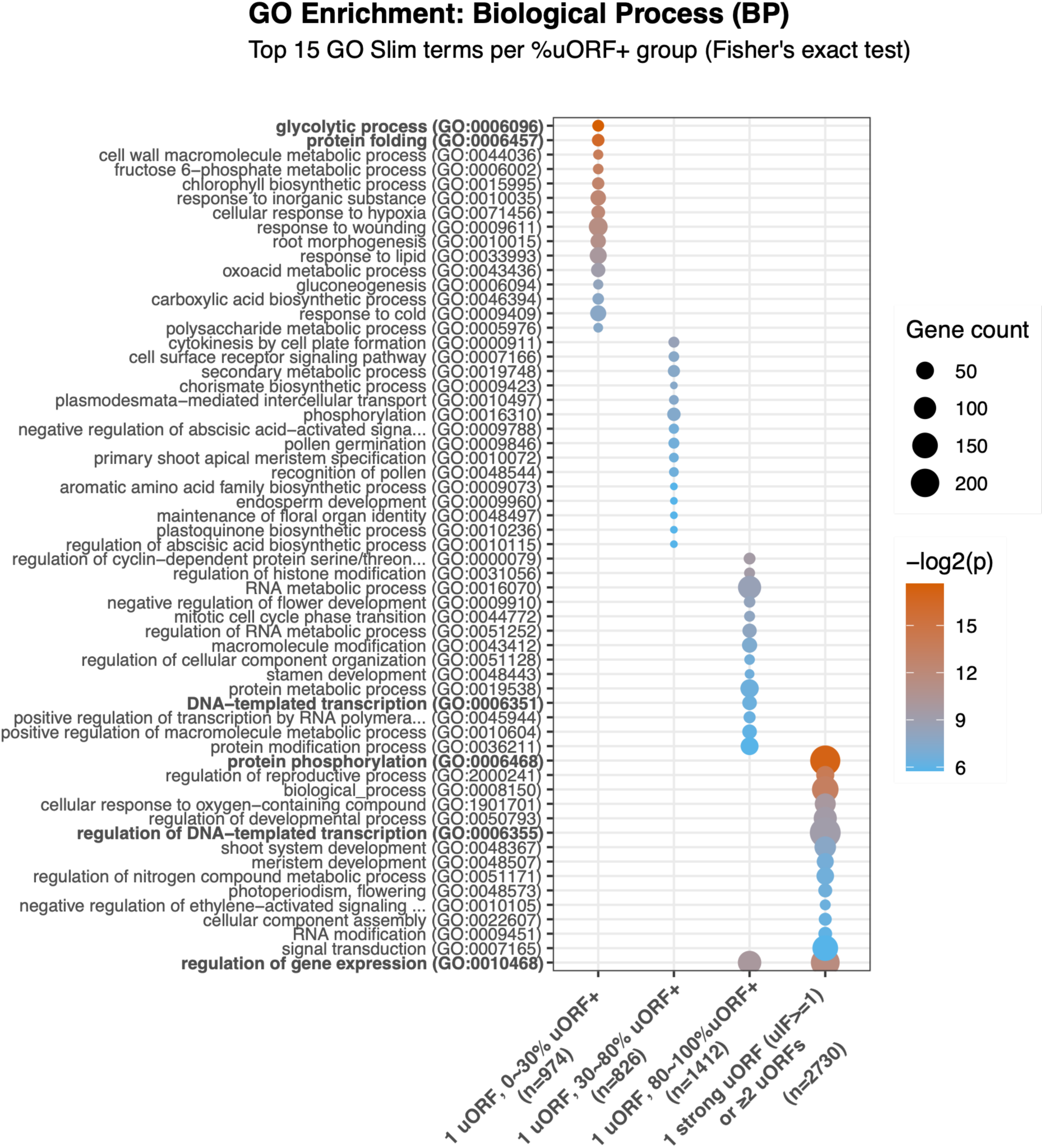
GO Biological Process terms enriched in uORF groups across different %uORF+ categories.

**Figure S9:**
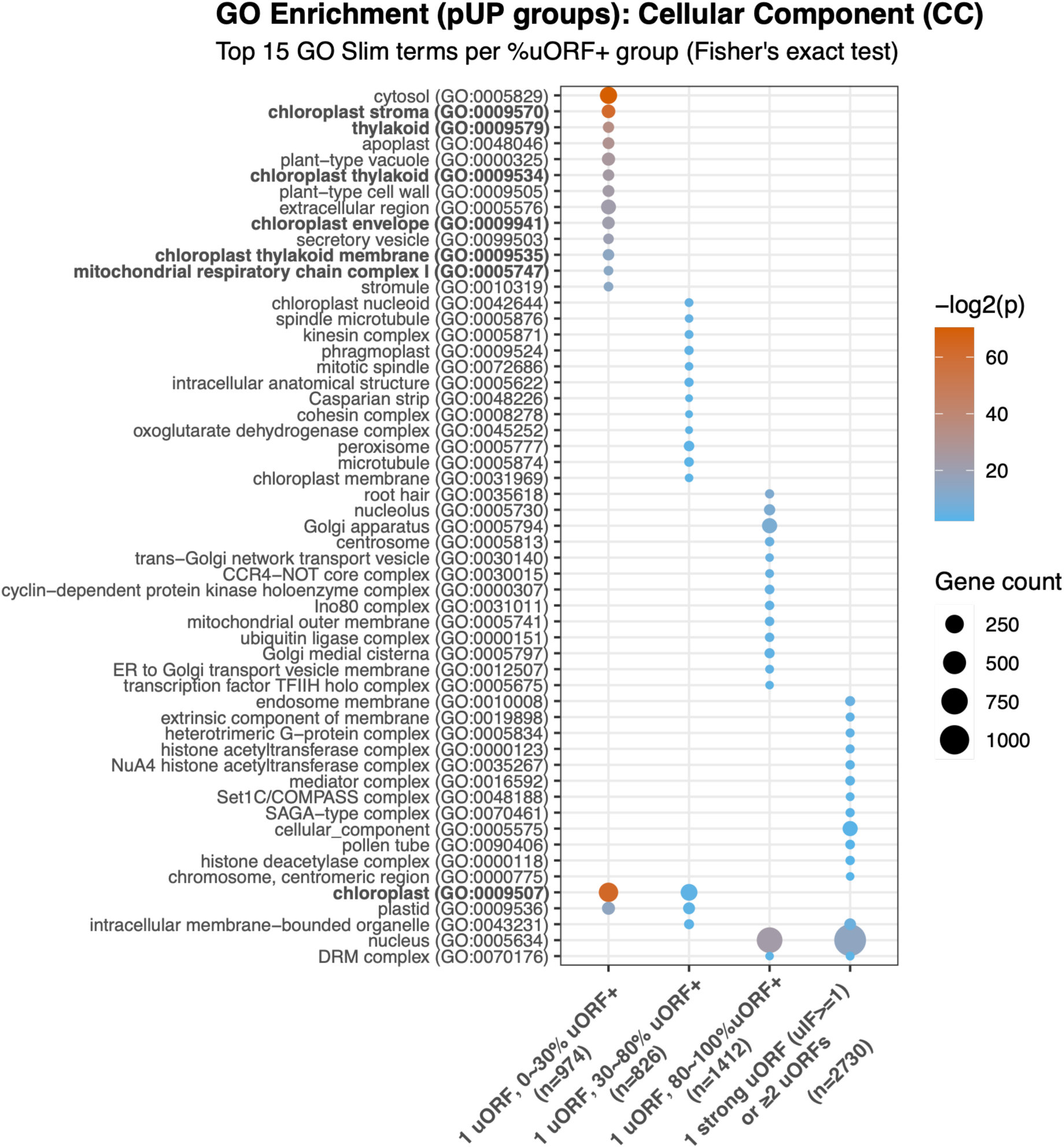
GO Cellular Component terms enriched in uORF groups across different %uORF+ categories.

